# Neuropathological hallmarks during the chronic phase of ischemic stroke in mice and humans

**DOI:** 10.1101/2025.04.16.649173

**Authors:** Romeesa Khan, Gary Guzman, Trang Do, Patrick Devlin, John Ahn, Chunfeng Tan, Venugopal Reddy Venna, Sean P. Marrelli, Haniyeh Koochak, Claudia M. Di Gesù, Anthony Flores, Louise D. McCullough, Rodney M. Ritzel

## Abstract

**Background:** Improvements in acute stroke treatment, including endovascular thrombectomy and critical care management, have increased survival rates post-stroke. However, stroke remains a leading cause of long-term disability and many survivors have significant neurological and cognitive deficits. Despite this, the chronic neurological sequelae and underlying secondary injury mechanisms induced by ischemic stroke remain understudied.

**Methods:** This study examined long-term neurobehavioral recovery and neuropathology at 2- and 6- months post-stroke in young (12-week-old) male C57Bl/6 mice after a 60-minute transient middle cerebral artery occlusion (MCAO) or sham surgery. Behavioral testing included the open field test (OFT), novel object recognition test (NORT), fear conditioning (FC), nesting activity, and tail suspension. Post-mortem brain samples from patients with chronic ischemic stroke were also assessed. Immunohistochemistry (IHC) was performed to assess demyelination (MBP), neuronal apoptosis (TUNEL), and Aβ42 in human brains. Flow cytometric analysis was performed to assess microglial phenotypes, the chronic neuroimmune landscape, and to evaluate senescent-like phenotypes (SA-βGal and lipofuscin). Transcriptomic profiling was performed using RNA isolated from the ipsilateral hemisphere in stroke mice.

**Results:** Experimental stroke caused progressive cognitive and motor decline up to 6 months post-MCAO. IHC and flow cytometric analyses revealed a significant increase in TUNEL-positive neurons, cortical and hippocampal gliosis, white matter degradation, senescent cell accumulation, and altered microglial function. IHC analysis of postmortem human brains shows significantly increased levels of microgliosis, senescent cells and amyloid burden. Transcriptomic analysis revealed that pathways involving apoptosis, microglial activation and the complement pathway were chronically upregulated after stroke.

**Conclusion:** Our findings demonstrate that ischemic stroke induces a non-resolving microglial response and accelerated inflamm-aging in the brain, evidenced by premature senescence and elevated production of cytokines within the chronic infarct microenvironment. Senescent-like phenotypes and chronic neurodegenerative disease signatures may contribute to the progressive worsening of cognitive function post-stroke. These results suggest that chronic, ongoing neurodegeneration occurs late after stroke, even in younger mice. Mitigating these detrimental changes may offer viable targets for delayed treatment strategies for stroke.

## Introduction

Stroke is the fifth leading cause of death in the United States and one of the leading causes of long-term disability^1^. Approximately 800,000 people experience a new or recurrent stroke each year, the vast majority of which are due to ischemic stroke^2^. Although survival and recovery post-stroke have improved due to advancements in acute stroke management and thrombolytic treatments, such as Alteplase, the long-term burden of post-stroke disability persists^3^. Approximately 40 percent of patients surviving an ischemic stroke have poor functional outcome and difficulties in living independent lives, even a decade after stroke^4^. In addition to motor disability, the long-term sequelae of ischemic stroke include depression, seizures, fatigue, cognitive impairment, and dementia. The risk of dementia is approximately doubled after a stroke, and is disproportionately higher in aged individuals^5^. The secondary injury mechanisms underlying the onset and progression of post-stroke cognitive decline remain unknown.

Approximately fifty percent of people who experience post-stroke cognitive impairment are diagnosed with vascular dementia, or vascular cognitive impairment (VCI), but data suggest that the development of dementia may be a product of both VCI and degenerative pathologies^6^. VCI is caused by multiple factors and can be attributed to persistent, or chronic, cerebral hypoperfusion caused by ischemia which leads to an altered brain environment^7^. One factor contributing to VCI is neuroinflammation, which triggers an immune response in the central nervous system (CNS) and recruitment of peripheral immune cells to the brain.

The resident immune cells of the brain, microglia, are among the first to respond to ischemia. Microglial activation is a hallmark of post-stroke pathology during the acute phase, but recent studies suggest these cells may affect neuroinflammatory processes long after the occurrence of a stroke. However, the vast majority of pre-clinical studies have only evaluated subacute (i.e., 30 days) outcomes in rodent models^8, 9^. Microglia display a continuum of activation phenotypes depending on environmental cues and can promote both protective and detrimental responses after stroke. The early microglial response has been linked to the regulation of astrocytic and neuronal function; however, a pronounced inflammatory phenotype emerges during the more chronic phase of brain injury^10^. At later stages, microglia appear to secrete inflammatory factors and adopt a de-ramified, hypertrophic appearance consistent with morphological activation^10^. Microglia exposed to the chronic inflammatory environment induced by ischemic stroke may develop a senescent-like phenotype and contribute to later progressive neurodegeneration.

Cellular senescence can be induced by the gradual increase in age-related inflammation termed ‘inflamm-aging’ via a positive feedback loop^11^. Chronic stress, inflammation, and oxidative damage are all factors contributing to cellular senescence, even in younger individuals^12^. Senescent cells adopt a senescence-associated secretory phenotype (SASP) which is characterized by enhanced secretion of inflammatory cytokines and other neurotoxic soluble molecules^13^. Recent studies show that biomarkers of cellular senescence are upregulated during the acute phase of ischemic stroke^14, 15^, however, the chronicity of these changes and the secondary mechanisms underlying cognitive dysfunction at late stages after stroke are currently unknown. The mechanisms governing post-stroke neuroinflammation, inflamm-aging, and cellular senescence are likely interconnected and may provide insights into the overall process of brain aging.

We hypothesize that ischemic stroke is a chronic neurodegenerative disease due to non-resolving inflammatory processes, including premature senescence, which persist and worsen chronically after stroke. To test our hypothesis, we subjected wild-type (WT) mice to transient middle cerebral artery occlusion (MCAO). We utilized behavioral tests, transcriptomic profiling, flow cytometry, and immunohistochemistry to assess neuropathological hallmarks at six months post-MCAO, which were confirmed in postmortem human brain samples. To determine if stroke *accelerates* age-related brain pathologies, we utilized younger mice which exhibit more resilient long-term functional outcomes to acquired brain injury than their older counterparts^16,17^.

## Methods

### Animals

Wildtype (WT) C57BL/6 male mice (12-weeks-old) were obtained from the Jackson Laboratory. All mice were received at least 6 weeks prior to any experimental manipulations. Animals were group-housed in Tecniplast individually ventilated cage (IVC) racks and had access to chow and water *ad libitum*. Rooms were maintained at 21–23°C, under a 12:12-hour light:dark cycle. Animal procedures were performed at an AAALAC accredited facility and were approved by the Animal Welfare Committee at the University of Texas Health Science Center at Houston, TX, USA.

### Transient middle cerebral artery occlusion (tMCAO)

Cerebral ischemia was induced under isoflurane anesthesia in 3-month-old male WT mice by occluding the right middle cerebral artery, as previously described^18^. Body temperature was maintained at 37°C throughout the surgery by a feedback-controlled heating pad and rectal temperature probe. A midline ventral neck incision was made and unilateral middle cerebral artery occlusion (MCAO) was performed by inserting a silicon-coated monofilament into the right internal carotid artery which was advanced into the circle of Willis and just beyond the origin of the MCA, thus blocking flow to the right MCA. Following one hour of ischemia, the monofilament was withdrawn to establish reperfusion. The animals were placed in a recovery cage with a heating pad, subcutaneously administered with 150 μl of saline and provided with wet mash for 7 days post-surgery. Mice were euthanized 6 months after MCAO. Sham controls underwent the same surgical procedure with the exception of monofilament insertion. Neurological deficit scoring (NDS) post-surgery was performed according to the following scoring paradigm (Bederson-score): 0, no deficit; 1, forelimb weakness and torso turning to the ipsilateral side; 2, circling to the affected side; 3, inability to bear weight on the affected side; and 4, no spontaneous locomotor activity. All analyses were performed by investigators blinded to surgical conditions. Animals were pre-assigned to either stroke or sham groups by computer randomization.

### Consideration of sex as a biological variable

It has been previously reported that estrogen is neuroprotective^19^, may induce post-stroke neurogenesis and contribute to post-stroke recovery in a mouse model of stroke^20^. As this study utilized young mice, we only included male mice to avoid these confounding effects in young female mice. The human post-mortem brain cohort includes both male and female patients, however, the female samples are all from post-menopausal (>65 years-old) individuals.

### Behavioral Assessment

All behavior tests were performed at baseline, prior to surgery. Post-surgery, a series of behavioral tests was performed monthly over a period of five days. The first three days were used to complete the Open Field and Novel Object Recognition tests. The nest building score was assessed on the fourth day, and tail suspension was performed on the fifth day. The fear condition test was only performed once, one week prior to the study endpoint.

### Open Field Test (OFT)

The OFT was used to assess spontaneous locomotor activity as well as anxiety-like behavior^21^. Mice were placed in the center of an open field chamber and total distance traveled was quantified in total centimeters traveled over a period of 10 minutes, tracked by the Ethovision system. Anxiety-like phenotype was measured as the total amount of time spent in the center of the arena compared to the border, with longer duration in the border region suggesting anxiety-like behavior. Each trial was 10 minutes long and data was calculated as a percentage of the total duration.

### Novel object recognition test (NORT)

The NORT is used for evaluation of cognitive function and learning memory in rodents^22^. Mice have an innate tendency to explore novel objects more than familiar, or old, ones. This test uses the discrimination index (DI) score to assess how much preference a mouse gives to the novel object compared to the familiar object, with a DI of 0 indicating no preference, a positive DI indicating preference towards the novel object, and a negative DI indicating preference for the familiar object. Animals were habituated to the test arena during OFT. 24 hours after open field, mice were placed in the familiar arena with two identical objects. Mice were allowed to explore for a total of 30 minutes maximum, with data being recorded for the first 30 seconds for object exploration. The next day, mice were placed in the same arena again with a novel object and a familiar object. The time spent exploring each object was recorded (EthovisionXT) and the DI was calculated by using the formula DI = (NT – ST) / (NT + ST), where NT is time spent exploring the novel object and ST is time spent exploring the same (familiar) object.

### Tail suspension test (TST)

The tail suspension test is a measure of depressive-like phenotypes^23^. Mice were suspended from TST apparatus consisting of an elevated metal rod placed horizontally. Each trial lasted 6 minutes and the recording was used to calculate total amount of immobility time as a percent of the total trial duration. Higher immobility time represents heightened depressive-like behavior.

### Nest building

Nest building was used to assess overall well-being and apathy. The test was performed and scored based on the Deacon protocol and rating system^24^. At the start of the dark cycle, each mouse was housed individually overnight in a new cage with normal food and water access. The next morning, nest constructions were scored on the qualitative scale and the untorn nestlets were weighed to quantify nesting activity. The data was recorded and displayed as the overall score per mouse (1-5).

### Fear conditioning

Fear conditioning was used to measure contextual memory, as previously described^25^. Briefly, mice were placed in a rectangular box and underwent two phases, training/habituation and shock, on day one. On day two, the mice underwent a third phase, memory testing. The amount of freezing time, or inactivity state, was recorded by EthoVisionXT tracking software. The first minute of the training and testing phases were used to analyze differences in freezing time, with decreased freezing behavior in the test phase indicating impaired memory function.

### qRT-PCR

Quantification of relative mRNA expression levels for cdkn2a (p16), Bcl2, cdkn1b (p27), and p53 was performed by extracting RNA from mouse brain hemisphere homogenates using the RNeasy mini kit (QIAGEN, Cat#74104). Then 500 ng of mRNA was reverse-transcribed to cDNA using the RevertAid First Strand cDNA Synthesis Kit (Thermo scientific, Cat#K1621). Reverse transcriptase real-time (RT) PCR was performed using the BioRad CFX384 or BioRad CFX96 systems. The RT-PCR reaction mix for the 96-well system (adjusted with dH2O to a total volume of 20 µl) contained 1 µl of sample cDNA, 10 µl TaqMan Fast advanced master mix (Thermo Fischer, USA), and 1 µl of the respective TaqMan probes (Cat#4331182) to genes cdkn2a (Mm00494449_m1), cdkn1b (Mm00438168_m1), Bcl2 (Mm00477631_m1), p53 (Mm01731290_g1) and GAPDH (Mm99999915_g1). Relative mRNA target gene expression levels were calculated according to the ΔΔCt method^26^ and were normalized to the housekeeping gene, GAPDH.

### Nanostring analysis

#### NanoString nCounter Analysis System

After extracting RNA from the samples, RNA quality was assessed using Agilent TapeStation, and the its quantity was measured using the Invitrogen™ Qubit™ RNA High Sensitivity (HS) kit. Subsequently, the samples underwent an overnight hybridization process at 65°C in a thermocycler. During this process, RNA molecules (analytes) were paired with nCounter optical barcodes (from the nCounter® Mouse NeuroInflammation and NeuroPath Panels by NanoString Technologies). Following hybridization, the nCounter Pro Analysis System immobilized and digitally counted RNA molecules that were hybridized to the nCounter barcodes. This constitutes ex situ digital counting of RNAs. For this aim, the hybridized samples should be prepared before processing to the data collection. This preparation was performed on nCounter Prep Station and was involved purification and immobilization of samples within a sample cartridge. Later, on the nCounter Digital Analyzer, hundreds of images were captured for each sample, resulting in hundreds of thousands of counts of target molecules per sample. Following internal image processing, the results were stored as RCC (Reporter Code Count) files containing raw RNA counts.

#### nSolver

The RCC raw data was imported from the Digital Analyzer into nSolver Analysis Software version 4.0 (NanoString Technologies) to go through normalization process. Standard normalization uses a combination of Positive Control Normalization (which uses synthetic positive control targets to adjust for variations that exist across samples, lanes, and cartridges), and CodeSet Content Normalization (which uses housekeeping genes to adjust for differences in analyte abundance and/or analyte quality across samples). Differential Gene Expression (DE), PathView, Gene Set Analysis (GSA), Cell Type Profiling, and Pathway Scoring were achieved using nSolver Advanced Analysis 2.0.134 Software. A P-value < 0.05 was considered statistically significant. The Benjamini-Yekutieli method was applied to adjust the p-value as the False Discovery Rate (FDR).

#### ROSALIND® Nanostring Gene Expression Analysis

Data was also analyzed by ROSALIND® (https://rosalind.bio/), with a HyperScale architecture developed by ROSALIND, Inc. (San Diego, CA, USA). Read Distribution percentages, violin plots, identity heat maps, and sample MDS plots were generated as part of the quality control (QC) step. Normalization, fold changes, and *p*-values were calculated using the criteria provided by <u>Nanostring</u>. ROSALIND follows the nCounter Advanced Analysis protocol for dividing counts within a lane by the geometric mean of the normalizer probes from the same lane.

Differential gene expression analysis was performed using the Student’s *t*-test on log2-transformed and normalized gene expression. *P*-values were adjusted using the Benjamini– Hochberg method to estimate false discovery rates (FDR).

### Flow cytometry

Flow cytometry sample preparation and staining were performed as reported previously^27^. Briefly, mice were trans-cardially perfused with 40 ml of cold sterile saline prior to removal of brain tissue. The ipsilateral brain hemisphere was isolated, the olfactory bulb and cerebellum were then removed, and the remaining hemisphere was filtered through a 70-μm filter into a 50 ml conical tube. Cells were resuspended in complete Roswell Park Memorial Institute (RPMI) 1640 (Invitrogen, Cat# 22,400,105) medium and digested in collagenase/dispase (1 mg/ml, Roche), papain (5 U/ml suspension, Worthington Biochemical), .5mM EDTA (Invitrogen, Cat#15575020), and DNAase I (10 mg/ml, Roche) for 45 minutes in a shaking incubator (200 rpm). The cell suspension was washed with 5 ml of RPMI, centrifuged at 500 g for 5 minutes at 4°C, then resuspended in a final volume of supplemented RPMI at 5 ml/hemisphere and kept on ice until staining. Cells were distributed into FACS tubes for staining and washed with 1 ml of FACS buffer prior to staining. The cells were incubated with Fc block (Anti-CD16/32, Biolegend, Cat#101302) for 10 minutes on ice prior to staining for the following surface antigens: CD45-eF450 (30-F11, eBioscience, Cat# 48–0451−82), CD11b-APCeF750 (M1/70, Biolegend, Cat#101262), Ly6C-AF700 (HK1.4, Biolegend, Cat#128024), and CD369/Clec7a-APC (RH1, Biolegend, Cat#144306). The fixable viability dye Zombie Aqua (Biolegend, Cat#423102) was used for live/dead discrimination. Cells were then washed in FACS buffer, fixed in 2% paraformaldehyde for 10 minutes, and washed once more prior to adding 500 µl FACS buffer.

Intracellular staining was performed using a fixation/permeabilization kit (BD Biosciences, Cat#554714) and the following antibodies: Lamp1-PerCPCy5.5 (1D4B, BioLegend, Cat# 121626). For intracellular cytokine staining, cells were incubated with a 1:1000 concentration of Brefeldin A (Biolegend, Cat# 420601) solution in RPMI, for 2 hours in a 37°C water bath. Cells were then resuspended in Fc block on ice, stained for surface antigens as above, and incubated in 100 μl of fixation/permeabilization solution (BD Biosciences, Cat#554714) for 20 minutes at 4°C. Cells were then washed with 750 μl of 1X permeabilization/wash buffer (BD Biosciences, Cat#554714) and resuspended in an intracellular antibody cocktail containing the cytokine antibody: TNF-PE-Cy7 (MP6-XT22, Biolegend, Cat#506324).

Lipid droplets were measured using BODIPY 493/503 (Invitrogen 1:500, Cat#D3922). SA-βGal activity was measured using the Senescence β-Galactosidase Activity Assay Kit (Cell Signaling Technology, Cat#35302S), per the manufacturer’s instructions.

Data were acquired on a Beckman-Coulter CytoFlex LX cytometer using CytExpert v2.6 software and then analyzed using FlowJo (Treestar Inc.). At least 5–10 million events were collected for each sample. CountBright Absolute Counting Beads (15 μl/sample, ThermoFisher, Cat# C36950) were used to estimate cell counts per the manufacturer’s instructions. Data were expressed as counts/hemisphere. Leukocytes were first gated using a splenocyte reference (SSC-A vs FSC-A). Singlets were gated (FSC-H vs FSC-W), and live cells were gated based on Zombie Aqua exclusion (SSC-A vs Zombie Aqua-Bv510). Resident microglia were identified as CD45^int^CD11b^+^Ly6C^-^, whereas peripheral leukocytes were distinguished as CD45^hi^CD11b^+^ myeloid cells and CD45^hi^CD11b^−^ lymphocytes. Cell type–matched fluorescence minus one (FMO) controls were used to determine the positivity of each antibody.

### Human sample selection

Postmortem human brain samples and embedded tissue was obtained from the BRAINS Human Biorepository and Histology Core. The demographic information for human samples is listed in **Supplementary Table 1.** Two regions (hippocampus and thalamus) from each case were selected for histopathological analyses. The cases with chronic stroke were chosen based on their clinical and final neuropathology reports, and a further gross view and microscopic view examination. The chronic infarct lesion, which was characterized by cystic cavity formation macroscopically and cystic cavity surrounded by a dense rim of reactive astrocytes (glial scars) microscopically. For the control cases, no significant neuropathological abnormalities were observed.

### Immunohistochemistry (IHC)

IHC for Formaldehyde-Fixed Paraffin-Embedded human brain samples and mouse brain tissue were performed as described previously^28^. Briefly, mouse brains were harvested post-transcardial perfusion with PBS then fixed in 4% paraformaldehyde solution for 24 hours, followed by washes of 70% and 100% ethanol and xylenes to be embedded in paraffin. After deparaffinization, human and mouse samples were subjected to heat-induced antigen retrieval using the EZ-Retriever ® IR System (BioGenex). The antigen retrieval solution used was a pH 6.0 citrate buffer (cat#) for 20 minutes (human samples) or 10 minutes (mouse samples) at 99°C. After antigen retrieval, the samples were blocked in blocking buffer (5% normal donkey serum, 1% Bovine serum albumin, and .2% TritonX-100 in PBS) for 2 hours at room temperature. Then, the samples were incubated with primary antibodies: (antibodies and cat#s and concentrations), overnight at 4°C. Then the samples were incubated with secondary antibodies (antibodies and cat# and concentrations) for 2 hours at room temperature. Slides were then washed in dH2O and cover slipped using DAPI mounting media (cat#).

### Imaging and quantification of histopathology

Imaging was performed using the Leica THUNDER Imager DMi8, with a 20x lens. For human samples, four images were taken from each region (thalamus and hippocampus) of each sample. Each sample obtained was a FFPE region of the same hemisphere. For mouse samples, the entire section (whole mouse brain) was imaged, and regions of interest such as hippocampus, decided a priori, were imaged further. The region of hippocampus analyzed was approximately 4mm posterior to Bregma. Images were analyzed using the ImageJ software by an investigator blinded to case conditions. All images were corrected for background noise and consistent fluorescence intensity. Quantities of signal-positive cells were obtained using individual regions-of-interest (ROIs) within respective channels, based on the secondary antibody used. The quantities of double-positive cells were measured using overlay images consisting of both antibody channels, and the counts were based on reselection of double-positive ROIs. Data are expressed as mean values ± SEM for mean fluorescence intensity, and for mouse samples each data point represents one mouse. For human samples, data are presented as 5 data points per region and sample.

### Quantitative assessment of brain injury by MicroCT imaging

Experimental mice were transcardiac perfused with 10mL (1x) PBS-heparin and then 10mL 4% PFA at rate of 2mL/min. After PFA, 1mL of Omnipaque (300 mg I/mL) was injected at rate of 2mL/min. The intact mouse was immediately transferred to the MicroCT scanner for imaging of the head. The mouse was positioned on its back and secured in the imaging cassette. The head was further secured with a section of folded paper towel above and below the head to reduce potential for motion artifact during imaging. Scanning was performed with a Bruker SkyScan1276 CMOS system. Imaging was performed at 60 kV source voltage and 200 uA source current. The detector was binned to 2048x2048 and each projection image resulted from an average of 3 images to achieve a final resolution of 18.6 microns. 3D reconstructions were performed with NRecon (Bruker, version 2.0.0.5) and subsequently converted to DICOMs using DicomCT. 3D image analysis was performed with ITK-SNAP 4.0.2 for segmentation and calculation of brain volumes.

### Analysis and statistical methods

Statistical analysis was performed using an unpaired parametric Student’s unpaired t-test, and two-way repeated measures ANOVA (for time factor interaction). The main effects were tested, followed by a post-hoc analysis with all related *p* values adjusted using Sidak’s method for multiple comparisons. Statistical correlations between the averaged behavioral variables and levels of expression for flow markers and immunohistochemical markers, were determined in each experimental scenario using the Spearman method. Statistical significance was considered at *p* < 0.05, and *\*p<0.05, **p<0.01, ***p<0.001, and ****p<0.0001* convention was used in the presented figures. All statistical analyses were performed using GraphPad Prism 7. Outliers were identified and excluded post-statistical analyses using the Grubb’s test.

## Results

### Experimental stroke causes progressive cognitive decline and persistent depressive-like behavior in young mice

Kaplan Maier analysis of survival rates showed that mortality peaked acutely after stroke, and occurred during the first week after surgery. The majority of stroke mice survived to 6-months post-MCAO (**Fig 1a-b**). Compared to sham controls, chronic stroke mice had significantly lower body weights at 3 and 6-months post stroke, and did not show the expected increase in weight over time (**Fig 1c**). In the open field test, both sham and stroke mice displayed deficits in locomotor function over time, with body speed and total distance traveled in the open field arena decreasing from 2 to 6-months post stroke. There were no significant differences between the locomotor function (distance traveled) between sham and stroke mice (**Fig 1d**). In the NORT, chronic stroke mice exhibited deficits in recognition memory compared to shams, which progressively worsened over time (**Fig 1e**). At six months post-surgery, chronic stroke mice displayed both a significantly heightened depressive-like phenotypes in the tail suspension test (**Fig 1f**) and impaired overall well-being in the nest building test (**Fig 1g**) compared to sham controls. Analysis of fear conditioning data also indicated significant impairment in cognitive function and learning memory in stroke mice compared to sham controls at six months after injury (**Fig 1h**).

**Figure 1.**
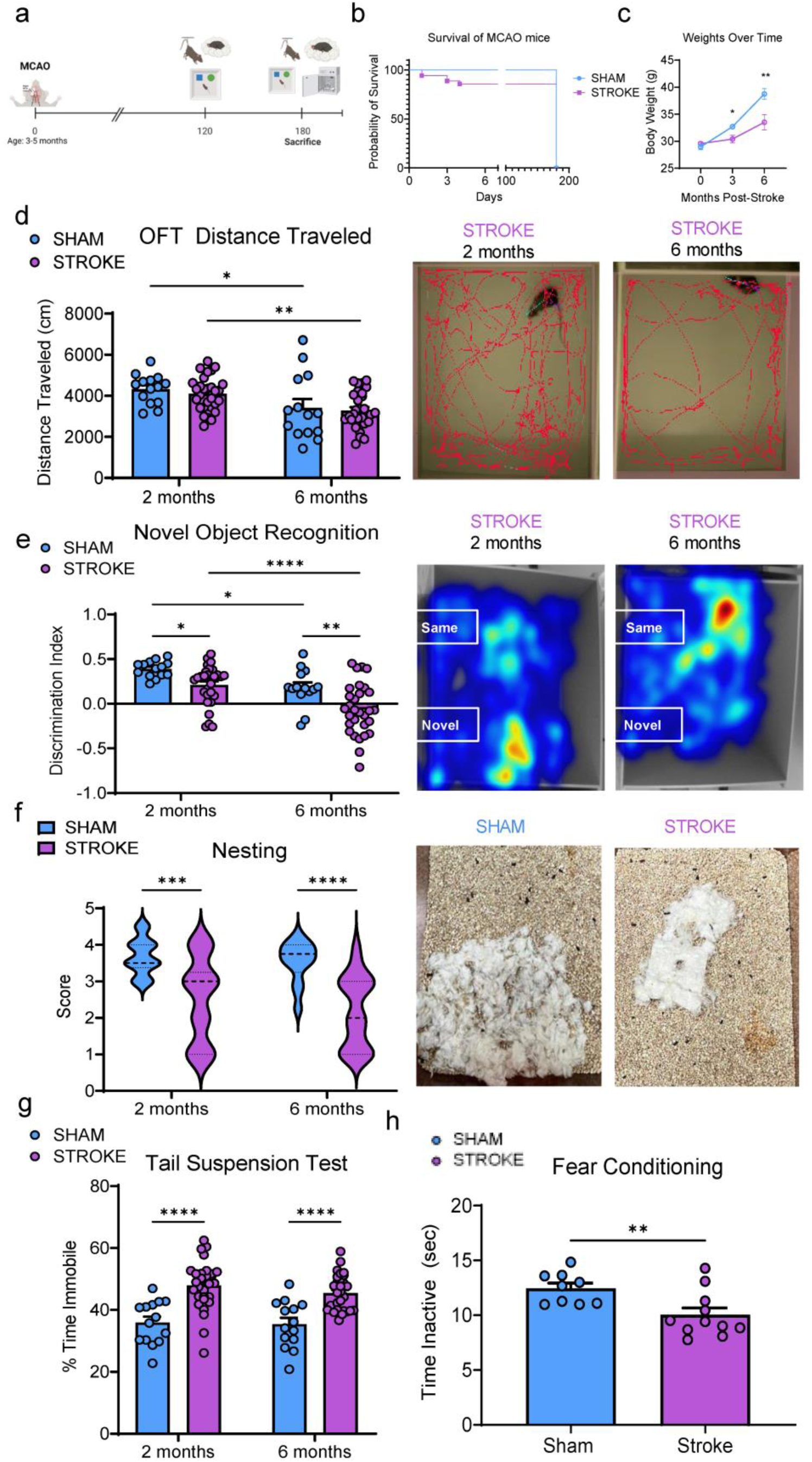
Cognitive dysfunction and depressive phenotypes persist chronically after stroke in mice. (**A**) Schematic showing a timeline of the experimental paradigm. (**B**) Kaplan-Meier plot of survival rate for MCAO and sham mice shows acute mortality in chronic stroke (cST) group which does not persist chronically. (**C**) Chronic stroke mice had significantly decreased body weights at 3 (p=0.0152) and 6-months (p=0.0050) post-stroke compared to shams. (**D**) cST mice had significantly decreased motor activity and speed at 6 months post stroke compared to 2 months post stroke in the open field test. Representative images from the arena tracking are shown, Two-way ANOVA Tukey’s multiple comparisons test, **p<0.01. (**E**) Compared to shams, cST mice had significantly increased immobility time in the tail suspension test compared to shams, indicating depressive phenotype, Two-way ANOVA Tukey’s multiple comparisons test, ****p<0.0001. (**F**) cST mice display significantly decreased nest-building performance compared to shams at both 2- and 6-months post-stroke indicating impairment of overall well-being. (**G**) In the NORT cST mice have significantly impaired cognitive function compared to shams, a phenotype that is exacerbated at 6 months post stroke, Two-way ANOVA Tukey’s multiple comparisons test, ****p<0.0001. (**H**) cST mice showed significant cognitive impairment in a fear conditioning test compared to shams (p=0.0084), unpaired t-test, p**<0.01.

### Transcriptomic profiling of stroke and sham mouse brains at six months post stroke

Transcriptomic analysis of the ipsilateral hemispheres at six months post-stroke revealed significant differences in gene expression between the stroke and sham groups. Heatmap visualization of pathway-specific gene signatures (**Fig 2a**) indicated that gene expression profiles within the stroke samples were highly correlated and distinct from the sham group, demonstrating differential pathway activation following stroke. Principal component analysis (PCA; **Fig 2b**) further supported these findings, with clear separation of stroke and sham samples in principal component space, highlighting distinct transcriptomic profiles between the two conditions. Volcano plot analysis (**Fig 2c**) identified numerous genes with significant differential expression between stroke and sham groups. Notably, Mmp12 was significantly upregulated in the stroke group (log2 fold change > 2.0; p < 0.05), along with other genes implicated in post-stroke recovery and senescence pathways. To validate these transcriptomic findings, qPCR analysis of Mmp12 expression was conducted (**Fig 2d**). The results confirmed significantly higher Mmp12 expression at 6 months in stroke brains compared to sham brains, consistent with the differential expression observed in the transcriptomic data. Further qPCR analysis of genes involved in senescence and apoptosis pathways demonstrated upregulation of p16, p21, p27, and Bcl2 in stroke samples compared to sham controls (**Fig. 2e**). Notably, p21 and Bcl2 were significantly upregulated (p < 0.05), while p27 showed a trend toward increased expression (p = 0.0761). These results suggest persistent activation of senescence and apoptotic pathways in the brain post-stroke.

**Figure 2.**
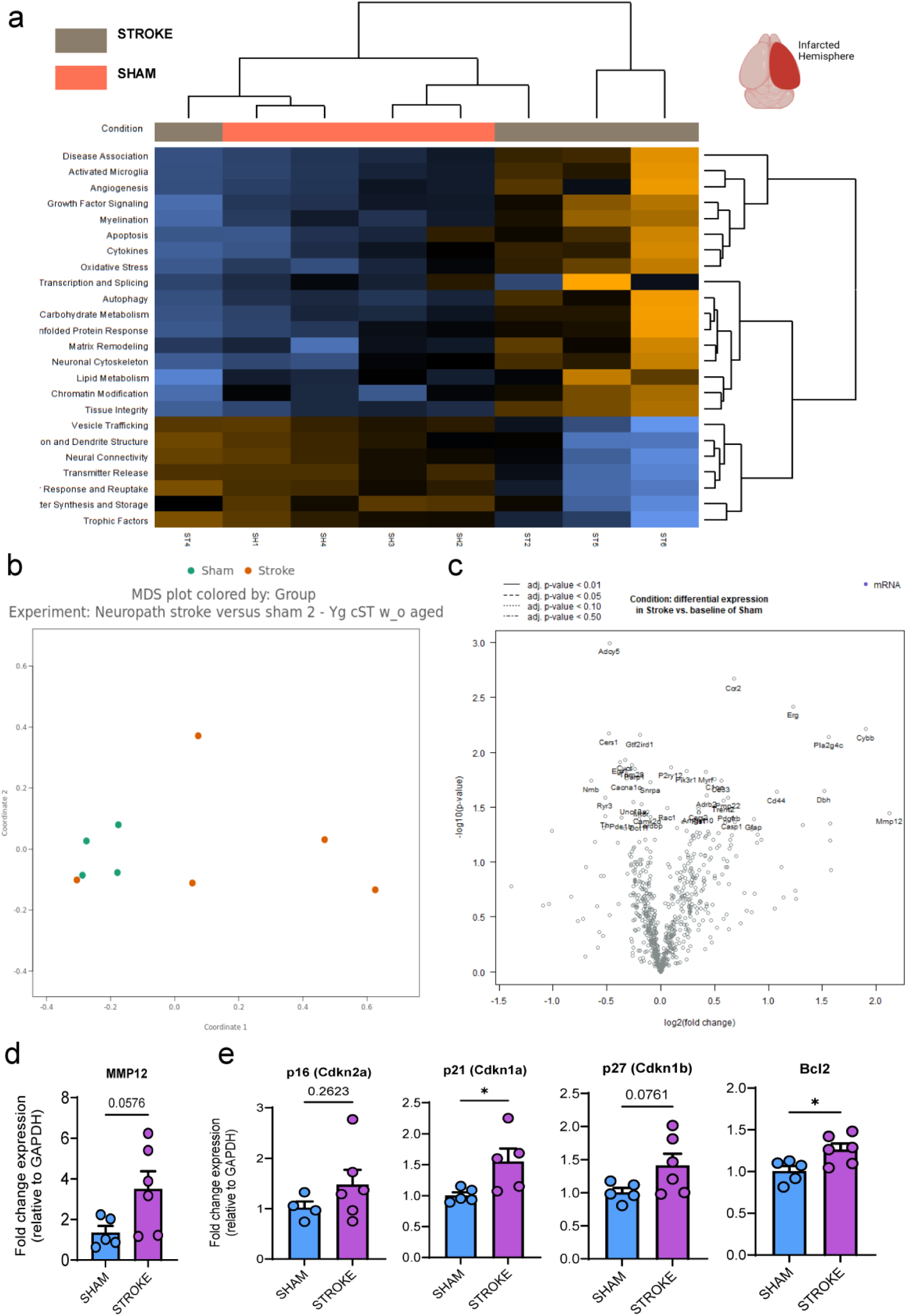
Transcriptomic analyses of ipsilateral mouse brain hemispheres at six months post stroke. (**A**) Heatmap visualization of pathway gene signatures as they correlate to condition (stroke versus sham). Gene and pathway signatures mostly correlate between stroke samples while the sham samples correlate together, indicating differential gene expression between the groups at six months post-stroke. (**B**) PCA of groups at 6 months post stroke. (**C**) Volcano plot of differential gene expression between stroke and sham. (**D**) qPCR gene expression analysis of relative MMP12 expression reveals significantly higher MMP12 gene expression in stroke brains compared to shams (p =.0056), serving as a validation of findings shown in the volcano plot and transcriptomic data. (**E**) qPCR analyses of various genes upregulated in senescence and apoptosis (p16, p21, p27, and Bcl2) pathways, show upregulated gene expression of all senescence-related genes in stroke brains compared to shams, at six months post stroke.

### Ischemic brain injury led to chronic gliosis, white matter atrophy, and cellular senescence at six months post-stroke

MicroCT imaging was performed to measure hemispheric brain volumes of sham and stroke mice. Decreased hemispheric volume was seen at six months post-stroke (**Fig 3a-b**). Subsequent analysis of the brains showed increased numbers of GFAP-positive astrocytes in white matter of stroke brains (**Fig 3g**). IHC analysis of the ipsilateral hemisphere from stroke mice brains showed significant increases in TUNEL-positive cells (**Fig 3e**) compared to sham controls, indicative of progressive neurodegeneration.

**Figure 3.**
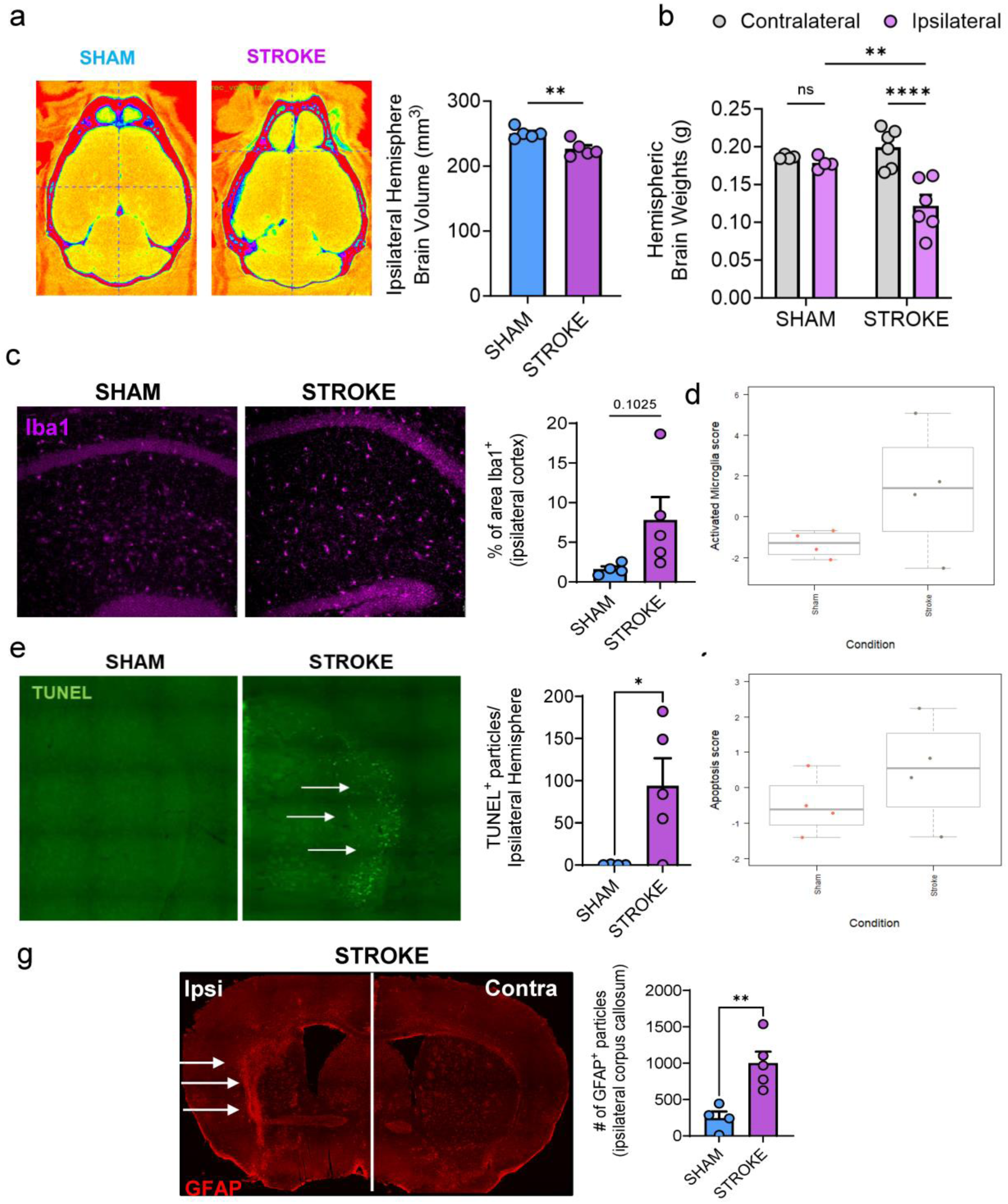
Histopathological analysis shows microglia are significantly altered in stroke mouse brains. (**A**) MicroCT representative images with quantification of brain volumes analyzed by MicroCT. Quantification of brain volumes (mm^3^) shows that sham brains have significantly higher volume of the ipsilateral (right) hemisphere compared to stroke brain, implying brain atrophy at 6 months most stroke (p = .0067, n = 5/grp). (**B**) Differences in right versus left brain hemispheres (ipsilateral versus contralateral) in sham and stroke brains show that there was no difference between the contralateral, or left, hemisphere in sham brains, however, the ipsilateral hemisphere weights of stroke brains were significantly less both between groups (p = .0027) and within the stroke group (p < .0001). (**C**) Representative images of immunofluorescent labeling for Iba1+ in mouse brain sections show increased microglial counts in the brains of cST mice. Quantification of Iba1^+^ signal in the ischemic hemisphere, ipsilateral cortex and ipsilateral corpus callosum of sham mice brains compared to chronic stroke (cST) mice. (**D**) Activated microglia pathway scoring from Nanostring analyses shows that the stroke group has significantly higher scores, or upregulated gene expression in microglial activation. (**E**) Representative images of TUNEL staining in the ipsilateral/right hemisphere of sham and stroke mouse brain. Quantification of TUNEL^+^ particles displays a significantly higher burden of TUNEL^+^ staining (green) in stroke brains compared to shams. (**F**) Transcriptomic analyses of upregulated gene signatures show a higher gene score for the apoptosis pathway signature in stroke brains. (**G**) Representative images of immunofluorescent labeling for GFAP^+^ in mouse brain sections show increased astrocyte signaling, and greater glial burden in the brains of cST mice in the ipsilateral hemisphere, compared to the contralateral hemisphere.

### Ischemic stroke induces chronic activation of microglia with a senescent-like phenotype

Our IHC findings demonstrated that Iba1-positive microglia (**Fig 3c**) display a significantly different morphology after MCAO compared to sham. Microgliosis appears to persist in the ipsilateral hemispheres, in both the cortex and hippocampal regions of stroke mice brains. Transcriptomic profiling of the ipsilateral brain hemispheres of stroke mice also showed increased genetic signatures of microglial activation genes and pathways in stroke mice but not in shams, shown as the Activated Microglia score by condition (**Fig 3d**).

### Ischemic stroke causes chronic alterations in microglial function, neuroinflammatory signaling, and oxidative stress

We next performed flow cytometric analysis of ipsilateral brain hemispheres in sham and stroke mice at six months post-injury. Compared to shams, the ipsilateral hemispheres weighed significantly less compared to the contralateral hemisphere (**Fig 3b**). After normalizing cell counts to tissue weight, we found significantly higher numbers of CD45^int^CD11b^+^Ly6C^-^ microglia in chronic stroke brains compared to sham controls (**Fig 4a**). Ischemic brain injury increased microglial TNF production (**Fig 4b**) and led to upregulation of the disease-associated microglia (DAM) marker, Clec7a (**Fig 4c**), indicating increased microglial-mediated inflammation. Consistent with our histological findings, we found that microglia had greater lysosomal burden and lipid accumulation chronically after stroke, as evidenced by increased Lamp1 expression (**Fig 4d**) and increased fluorescent intensity of BODIPY^493/503^ (**Fig 4e**). SA-βGal levels were also markedly increased in microglia at six months post-stroke, suggesting microglia may be important drivers of inflamm-aging and seno-inflammation in the chronic phase of ischemia (**Fig 4f**). Similar changes in function were found in other cell types, such as endothelial cells (**Supplementary Figures 1 and 2**).

**Figure 4.**
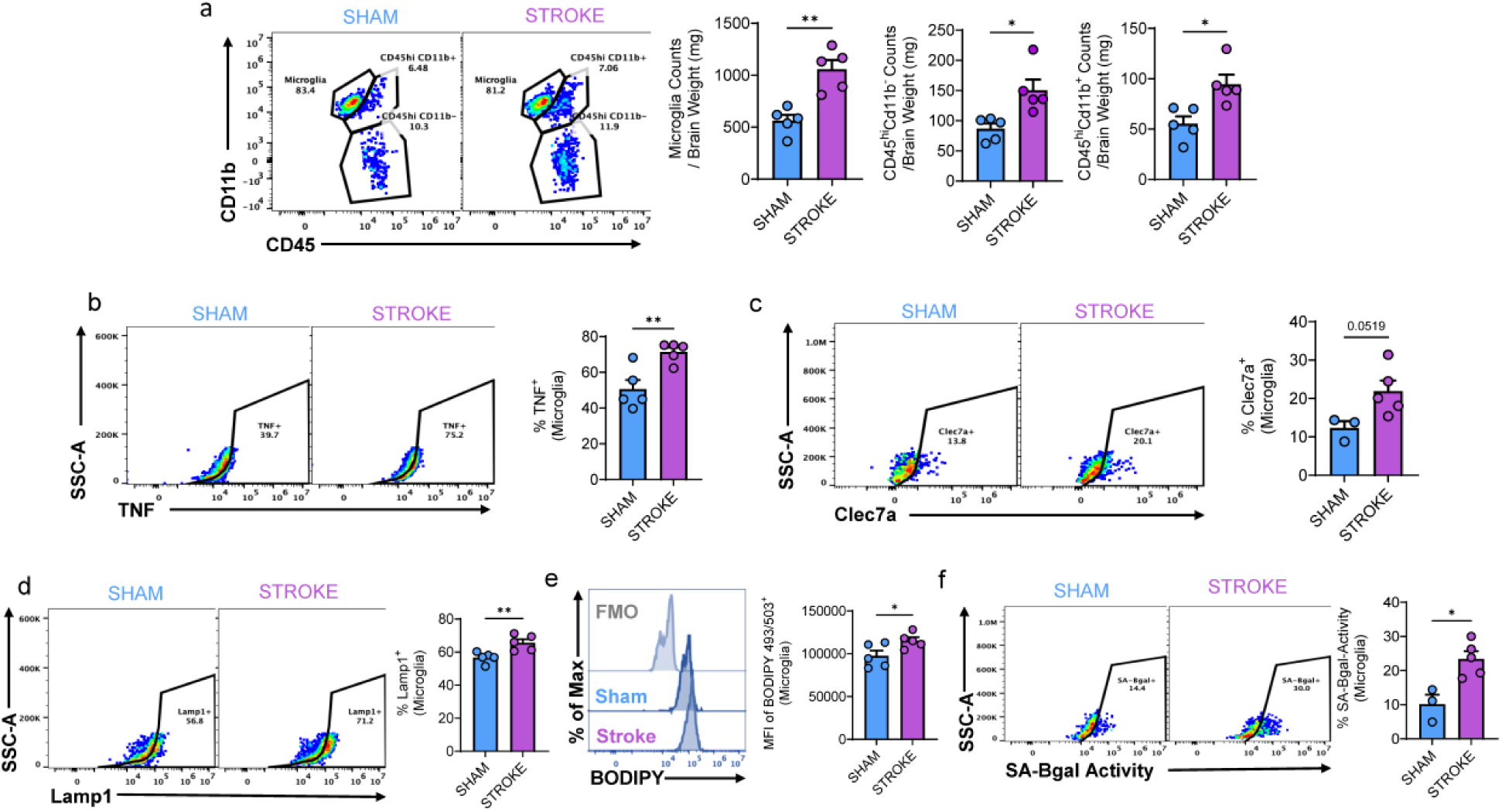
Functional alterations in microglia, neuroinflammation, and oxidative stress persist up to six months post MCAO. (**A**) Microglia counts (CD45^hi^CD11b^+^Ly6C^-^) normalized to hemisphere weight are significantly increased in stroke versus sham ipsilateral hemispheres (p=.0016). (**B**) Microglial production of TNF is higher in stroke brain ipsilateral hemispheres compared to shams (p=.0065). (**C**) Microglial Clec7a expression is increased in stroke brains (p=.0519). (**D**) Lamp 1 expression is higher in microglia of stroke brains compared to shams (p=.0081). (**E**) Fluorescence intensity of intracellular microglial BODIPY (p=.0409). (**F**) SA-βGal activity in microglia is significantly increased in stroke hemispheres compared to sham counterparts at 6 months post MCAO (p=.0105). N=3-5/grp.

### Cellular senescence, glial function alteration, and amyloid pathology found in postmortem human brain samples with chronic infarcts

IHC analyses of postmortem human brain samples from patients with chronic stroke showed persistence of many the same neuroinflammatory pathologies seen in mice with chronic stroke. The tissue sections assessed were located distal to the infarct, which implies the effects of stroke reach beyond the infarcted region (**Fig 5a**). Significantly increased levels of gliosis (GFAP^+^ astrocytes and Iba1^+^ microglia), SenTraGor^+^ senescent cells, and TUNEL-positive neurons (**Fig 5b-d**) were present in the hippocampal region of stroke brains compared to age-matched control brains. Fluorescence intensity of myelin staining was significantly decreased in hippocampi in stroke brains compared to age-matched controls, indicating that white matter loss and significant demyelination persists long after stroke (**Fig 5e**). Quantification of anti-Aβ42 staining in human brain samples showed evidence of increased amyloid plaque burden in the hippocampus of stroke samples, but not in controls (**Fig 5f**). Taken together, these findings suggest that prior history of ischemic stroke is positively associated with secondary neurodegenerative disease pathology in the hippocampus, independent of primary infarct site.

**Figure 5.**
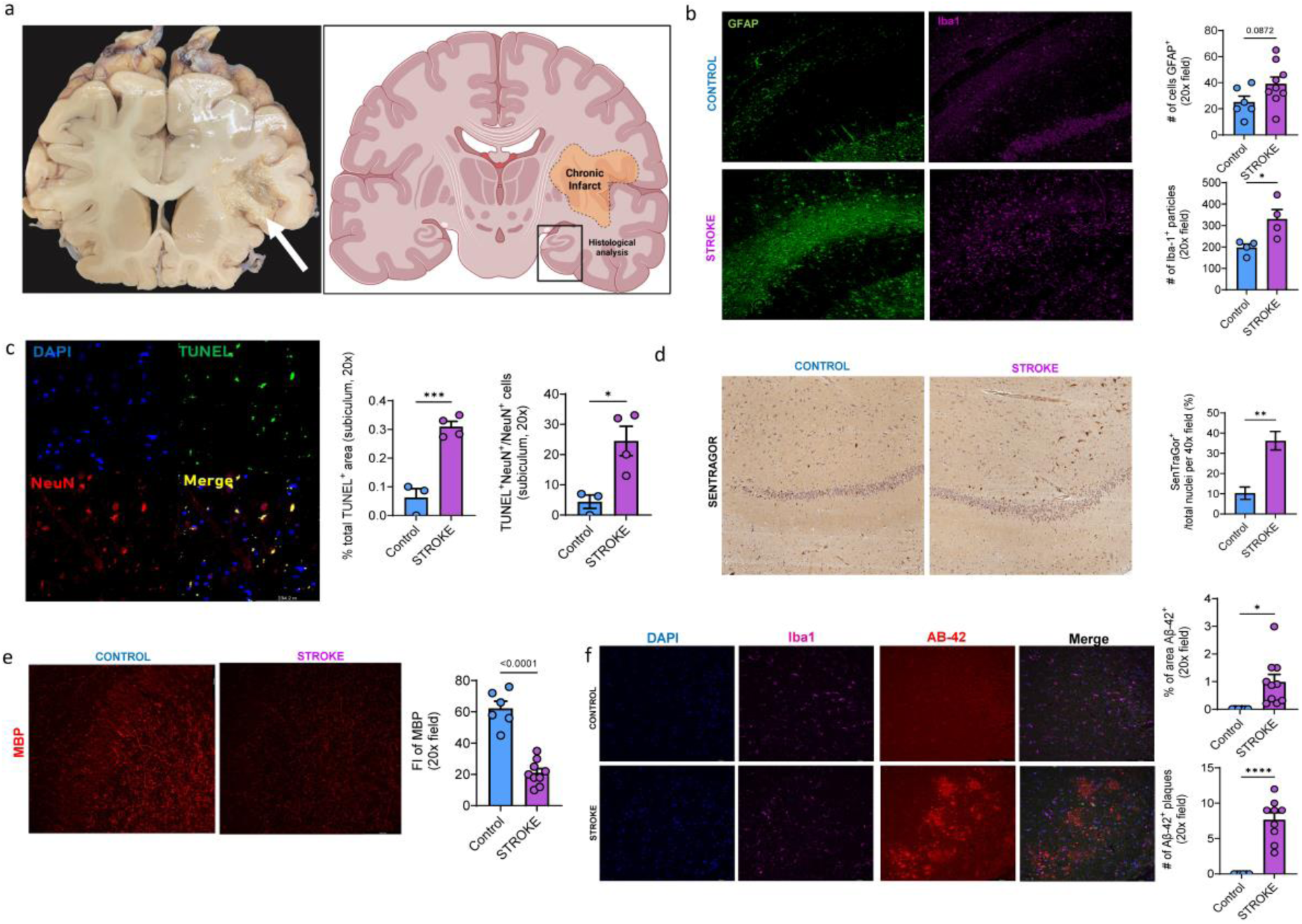
Cellular senescence, glial function alteration, and amyloid pathology found in postmortem human brain samples with chronic infarcts. (**A**) Gross histological image of human brain with chronic infarct. (**B**) Staining of GFAP and Iba1 in control and stroke samples of postmortem human brain hippocampus shows significantly increased glial counts in stroke cases compared to controls. (**C**) TUNEL co-labeling with NeuN-positive cells shows increased levels of neuronal apoptosis in stroke samples. (**D**) SenTraGor staining for lipofuscin indicates higher lipofuscin burden in the hippocampus of stroke samples compared to controls, indicating higher senescence burden chronically post-stroke. (**E**) Myelin Basic Protein staining in human stroke brains shows significantly higher MBP fluorescence intensity in stroke maples compared to controls, indicating degradation of myelin in stroke brains. (**F**) Images of DAPI, Iba1, and anti-Aβ42 staining in human brain samples. Higher AB-42 burden in stroke samples was indicated by increased levels of staining in stroke samples compared to controls, which displayed minimal amyloid-positive staining.

## Discussion

Our study yields several new findings. To the best of our knowledge, this is the first study to examine the neuropathological hallmarks and neurobehavioral sequela in mice at an extended timepoint, 6 months post-injury. This study offers insights into the dynamic, long-term effects of stroke on brain aging and neurological function. Our findings illustrate that stroke, even in younger animals with greater physiological resilience, precipitates a trajectory of progressive cognitive decline, neurodegeneration, premature senescent cell formation, and chronic microglial activation.

We observed that ischemic stroke leads to significant long-term cognitive and neurobehavioral deficits. Young mice subjected to tMCAO exhibited progressive cognitive impairments, including deficits in recognition memory. We also found significantly higher depressive-like phenotypes in mice that had a stroke compared to shams, although there was no effect of time on the severity of this phenotype. These results align with previous studies indicating that stroke survivors are at higher risk for enduring cognitive and emotional difficulties and poorer quality of life^29^.

Our findings reveal that ischemic stroke induces non-resolving brain inflammation and premature cellular senescence. Histological analyses demonstrated ongoing activation of microglia with a senescent-like phenotype, characterized by increased expression of p21 and accumulation of lipid droplets, especially in the cortex and hippocampal regions of the brain. This is consistent with the concept of ‘inflamm-aging,’ which describes the gradual increase in basal inflammatory states during the aging process^30^. We posit that ischemic stroke may accelerate or exacerbate brain inflamm-aging. The presence of senescent-like microglia, marked by increased SA-β-galactosidase activity, cytokine production, and lipofuscin/lipid accumulation, suggests that these cells may contribute significantly to chronic neuroinflammation and cognitive impairment. Our flow cytometry and IHC results show that these microglia are not only persistent but also exhibit enduring changes in their functional states, including increased TNF production and lysosomal burden, pointing to the presence of a chronic inflammatory environment in the post-stroke brain. The continued activation of these inflammatory networks is further evidenced by our transcriptomic data which shows chronic upregulation of most inflammatory pathways and gene signatures. These data align with previous studies that show similar findings in rodent models of stroke, albeit at less chronic timepoints post stroke^31, 32^.

The continued activation of these inflammatory networks is further evidenced by our transcriptomic data, which shows chronic upregulation of multiple inflammatory pathways and gene signatures. Transcriptomic analyses of the ipsilateral hemispheres at six months post-stroke revealed persistent activation of genes and pathways associated with inflammation, senescence, and neurodegeneration, consistent with previous reports^33^. We found increases in *P2RY12* and *TREM2* expression, both of which are genes associated with microglial activation and microglia-mediated neuroinflammatory processes^34, 35^. We also found that *MMP12* and *Cybb* expression is chronically upregulated post-stroke. *Cybb* expression has been linked to neuronal cell loss^36^, and may further indicate and validate our findings on neuronal apoptosis in human samples. Importantly, these results align with recent findings that chronic inflammation and cellular senescence play pivotal roles in post-stroke dementia and neurodegeneration^37^. These transcriptomic changes provide a deeper molecular context to the behavioral and histological phenotypes observed in our study, further underscoring the long-term impact of ischemic stroke on brain aging and pathology.

The observed white matter damage and increased gliosis in stroke mice, alongside findings of decreased myelin basic protein and elevated GFAP+ immunoreactivity, highlight the extent of chronic neurodegeneration. Importantly, our study also identifies an association between ischemic stroke and amyloid-beta plaque accumulation in postmortem human brain samples, indicating that stroke may accelerate or exacerbate Alzheimer’s disease-like pathology. However, it is important to note that the timing of onset for amyloid pathology, or the development of neurodegenerative disease is unclear. It has previously been shown that amyloid angiopathy elevates the risk for stroke^36^. However, due to the gap in understanding on the dynamics of amyloid pathology development, further investigation to understand whether ischemic stroke accelerates existing amyloid pathology or initiates its development, is necessary. This finding suggests a potential mechanistic link between stroke-induced neuroinflammation, cellular senescence, and amyloid pathology, which could be crucial for understanding the increased risk of dementia following stroke. This is consistent with prior reports of long-term structural changes and chronic inflammation following ischemic stroke^38^.

The evidence of persistent neuroinflammatory and neurodegenerative changes in human postmortem brain samples reinforces the translational relevance of our animal findings. The presence of higher senescent cell burden, phagocytic microglia, white matter loss, progressive neuronal apoptosis, and amyloid plaque formation in brains of stroke patients who had infarcts years earlier underscores the long-lasting impact of stroke on brain pathology. Importantly, all chronic infarcts in this sample set were located in regions distal to the hippocampus, highlighting the far-reaching effects of ischemic stroke. This suggests that therapeutic strategies aimed at modulating neuroinflammation and cellular senescence could be beneficial in managing long-term post-stroke disability and preventing the onset of dementia.

There are several study limitations to consider. The exclusion of female mice from the study does not allow us to investigate the specific effects of biological sex in experimental stroke models^39^. Additionally, the use of young mice limits our contextual understanding of the role of aging, the setting in which most clinical stroke naturally occur. While we maintain that the progressive cognitive decline and persistence of secondary injury mechanisms seen in younger, more resilient mice, are proof-of-principle that stroke accelerates brain aging, future studies must establish a more translational cohort of chronic stroke experiments by including both female and older mice to better model an at-risk population for stroke and post-stroke cognitive decline. Additionally, we recognize that the lack of longitudinal neuroimaging and histopathology hinders us from tracking the dynamics of accumulating senescent cell burden and chronic pathology post stroke.

In summation, our study provides evidence for the long-term and far-reaching effects of ischemic stroke. A deeper understanding of the chronic and persistent changes that occur post-stroke will create avenues for better insight into the molecular mechanisms that are chronically activated after ischemic brain injury. This information could aid in the development of novel delayed stroke therapeutic strategies. Future research will focus on elucidating microglial activation and neuroinflammation as major driver mechanisms of post-stroke cognitive impairment. Additionally, future studies will include a more translational cohort of middle-aged mice to better model the populations most at risk for chronic stroke outcomes.

## Acknowledgments

RK conducted all the experiments, carried out the statistical analyses, and wrote the manuscript. TD, GG, PD, JA, SM, HK, CDG, AF, and CT assisted with the experiments, figures, and samples. VV performed the surgeries. RR conceived and designed the project. RR and LDM provided guidance for the experiments and edited the manuscript. Schematics were made using BioRender.

---

**Supplementary Table 1.** Demographic and clinical characteristics of chronic stroke and control cases for postmortem brain samples.

| Case | Race | AGE | SEX<br>M/F | HTN | DM | Brain Weight<br>(gram) | Cause of Death | Location of the chronic<br>infarct lesion |
| --- | --- | --- | --- | --- | --- | --- | --- | --- |
| **Chronic stroke 1** | Caucasian | 68 | F | YES | YES | 1040 | Respiratory failure, ventricular fibrillation and cardiac arrest | Cerebellum and Midbrain (cerebral peduncle) |
| **Chronic stroke 2** | Hispanic | 66 | M | N/A | N/A | N/A | N/A | Right parietal lobe |
| **Chronic stroke 3** | Hispanic | 74 | F | YES | YES | 1130 | Heart failure, acute respiratory distress syndrome, coagulopathy and hemorrhagic shock | Bilateral basal ganglia and Left occipital lobe |
| **Chronic stroke 4** | AA | 67 | F | YES | YES | 960 | Diffuse alveolar damage, bronchopneumonia, bronchitis and bronchiolitis | Right temporal lobe |
| **Chronic stroke 5** | AA | 67 | F | YES | YES | 1120 | Multiorgan failure | Left frontal lobe |
|  |  | 68.4 | 1M:4F |  |  | 1062 |  |  |
| **Control 1** | AA | 65 | F | YES | NO | 1000 | Cardiac arrest |  |
| **Control 2** | Hispanic | 81 | M | YES | YES | 1040 | Hypoxemia |  |
| **Control 3** | AA | 74 | F | YES | YES | 930 | Acute hypoxemic respiratory failure with pulmonary thromboemboli |  |
| **Control 4** | AA | 97 | F | YES | YES | 980 | Acute myocardial infarction |  |
| **Control 5** | Hispanic | 62 | M | YES | YES | 1390 | Hypovolemic shock after splenic rupture |  |

